# Differential Transcript Usage Reveals Isoform-Level Remodeling of Tumor Biology in Clear Cell Renal Cell Carcinoma

**DOI:** 10.64898/2026.05.27.728189

**Authors:** Chinaza F. Nnam, Yiping Li, Minghui Zhang, Erick A. Mboya, Fred Kolling, Laurent Perreard, Thomas J. Palys, Elizabeth Pflugradt, Patricia A. Pioli, Marc Ernstoff, John D. Seigne, Jason R. Pettus, Bing Ren, Li Song, James Brugarolas, Brock C. Christensen, Lucas A. Salas

## Abstract

Clear cell renal cell carcinoma (ccRCC) is characterized by transcriptional reprogramming driven by hypoxia signaling, metabolic rewiring, and immune modulation. While gene-level analyses have defined key features of ccRCC biology, they do not capture isoform-level variation arising from alternative splicing. Differential transcript usage (DTU) represents an additional regulatory layer that may influence protein function, pathway activity, and clinical outcomes, yet its role in ccRCC biology and prognosis remains incompletely understood. We assessed differential expression in 127 ccRCC tumors and 33 normal-adjacent tissues from the Dartmouth Cancer Center cohort, with external validation in 94 CPTAC tumors, adjusting for cell-type proportions. DTU was identified using DRIMSeq/stageR, followed by limmavoom modeling with clinical and tumor microenvironment covariates. Transcript-based consensus clustering defined tumor subgroups, and Cox proportional hazards modeling integrated transcript-level features with clinical variables. In tumor versus normal comparisons, 1,170 transcripts exhibited significant differential usage, mapping to canonical ccRCC pathways with distinct patterns across functional and non-functional transcript classes. Consensus clustering based on transcript usage identified two subgroups with distinct angiogenic profiles and significant survival differences. Cluster-level analysis revealed DTU in genes involved in cytoskeletal organization (*ACTB*), immune processes (*B2M*), extracellular matrix organization (*FN1*, *APLP2*), and iron metabolism (*FTH1*) with protein domain alterations, including the loss of actin-associated domains in *ACTB* and immunoglobulin-like domains in *B2M*. Prognostic modeling identified twelve transcripts consistently retained across bootstraps, improving risk stratification over clinical variables alone. External validation confirmed overlapping prognostic transcripts, including *FGFR1* and *NUCB1*. Isoform-level features may serve as biomarkers and therapeutic targets in ccRCC.

**Statement of significance:** Transcript-level analysis uncovers potential regulatory pathways in ccRCC missed by gene-level approaches, revealing isoform-specific alterations that define survival sub-groups and offer potential biomarkers and therapeutic targets.

## 1. Introduction

Renal cell carcinoma (RCC) accounts for approximately 3% of adult malignancies and remains one of the ten most common cancers worldwide^1^. Clear-cell renal cell carcinoma (ccRCC) represents the predominant histologic subtype, comprising approximately 75% of newly diagnosed cases^2^. At the molecular level, ccRCC is defined by inactivation of the von Hippel–Lindau (*VHL*) tumor suppressor gene, resulting in stabilization of hypoxia-inducible factors and activation of a pseudohypoxic transcriptional program^3,4^. In addition, recurrent mutations in chromatin-modifying genes, including *PBRM1, SETD2, BAP1, and KDM5C*, occur in a substantial proportion of tumors, highlighting the central role of epigenetic dysregulation in disease pathogenesis^5,6^. Although targeted therapies directed at VEGF and mTOR signaling, as well as immune checkpoint inhibitors, have improved outcomes for patients with advanced disease, clinical behavior remains heterogeneous^7,8^. Current staging systems and clinicopathologic variables incompletely capture the biological heterogeneity of ccRCC and remain limited in their ability to predict disease progression and therapeutic response^9^. Molecular stratification has improved biological and clinical interpretation, including associations between alterations such as *PBRM1* mutation and favorable treatment response^10,11^. However, additional molecular insights are still needed to improve prognostication and patient stratification.

While transcriptomic profiling has significantly advanced our understanding of ccRCC, most studies have focused on gene-level expression changes^12,13^. However, alternative mRNA splicing, a process affecting more than 90% of multi-exon human genes, generates multiple transcript isoforms from a single gene and represents a major source of proteomic diversity^14^. In cancer, dysregulated splicing can produce aberrant splice variants that alter protein structure, localization, stability, and function, thereby contributing to oncogenesis independently of canonical gene-level alterations^15,16^. A well-known example is the AR-V7 splice variant in prostate cancer, which confers resistance to androgen receptor–targeted therapy^17^.

In ccRCC, emerging evidence suggests that transcript-level alterations may play an important role. Mutations in chromatin-modifying genes, widespread DNA methylation, and even retroelement dysregulation are known to influence splice-site selection and transcript regulation^18,19^. Prior studies have identified differentially expressed splice variants in ccRCC, including isoforms of *EGFR, SLC15A4, FGD1, SYNPO,* and *RNASET2*, and have demonstrated that splice-variant–based signatures may stratify patient survival across independent cohorts. Notably, the *SLC15A4* splice variant (locus 12:129299615–129307792) has been linked to hypermethylated ccRCC and *SETD2* alteration, connecting aberrant isoform usage to an epigenetic dysregulation that characterizes this disease^20^. Given the established roles of *SETD2* in H3K36 trimethylation, transcriptional elongation, and RNA splicing, these observations support the premise that transcript-level analysis may capture biologically meaningful variation not evident at the gene level^22,23^. Despite these advances, several critical gaps remain. First, many prior studies have focused on differential isoform expression, which measures absolute transcript abundance, rather than on formal modeling of differential transcript usage (DTU), which specifically evaluates shifts in the relative proportions of isoforms within a gene^24^. Second, tumor transcriptomes are confounded by variable proportions of immune, stromal, and angiogenic cells, making it difficult to disentangle tumor-intrinsic splicing alterations from microenvironment-driven signals^25^. Third, the extent to which molecular features, including DTU-derived features, provide independent and reproducible prognostic value beyond established clinical variables has not been comprehensively evaluated in ccRCC^26,27^. Systematic interrogation of DTU in ccRCC, therefore, offers an opportunity to refine our understanding of tumor biology at higher resolution. By modeling relative transcript usage while accounting for cellular composition, it is possible to identify isoform switching events, characterize coding and non-coding transcript dynamics, and define transcript-level signatures with potential clinical relevance.

In this study, we analyzed a cohort of 127 ccRCC patients from the Dartmouth Cancer Center Renal Tumor Bank using robust, covariate-adjusted differential gene expression and differential transcript usage analyses. Finally, we sought preliminary external validation of transcript-level prognostic features using the CPTAC ccRCC cohort. Our objectives were to define the landscape of isoform-level dysregulation in ccRCC and to determine whether transcript usage patterns provide prognostic information beyond established clinical variables. Through this approach, we aim to establish differential transcript usage as a biologically and clinically relevant dimension of ccRCC heterogeneity.

## 2. Study Design and Cohorts

### 2.1. Dartmouth Cancer Center cohort

The primary discovery cohort consisted of 160 clear-cell renal cell carcinoma (ccRCC) samples. from the Dartmouth Cancer Center (DCC) Renal Tumor Bank, including 127 primary tumors (109 with complete information) and 33 paired normal-adjacent kidney tissues. Samples were collected under institutional review board approval (IRB# STUDY02000672) between 1994 and 2014. Nephrectomy specimens were snap-frozen in liquid nitrogen immediately after resection and stored at −140°C until processing. Clinical and demographic variables, including age at diagnosis, sex, TNM stage, Fuhrman grade, and survival outcomes, were abstracted from the electronic medical record and curated into an analysis-ready phenotype table. For external validation, the Clinical Proteomic Tumor Analysis Consortium (CPTAC) ccRCC cohort was used. Isoform expression data (log2(FPKM)) were retrieved from https://www.linkedomics.org/data_download/CPTAC-pancan-CCRCC/. A total of 94 samples with complete clinical and expression data were included in the validation analysis.

### 2.2. DNA, RNA Extraction, Library Preparation, and Sequencing

DNA was extracted via Qiagen Blood and Tissue Kit using the tissue method. Then, the DNA samples were converted using the Tecan True-Methyl OxBS module according to the manufacturer’s instructions. Total RNA was extracted from frozen tissue using the Qiagen AllPrep DNA/RNA mini kit according to the manufacturer’s instructions. RNA integrity was assessed using an Agilent 2100 Bioanalyzer, and only samples with RNA integrity number (RIN) ≥ 5 were advanced to library preparation. Ribo-depleted libraries were generated using the Illumina TruSeq Stranded mRNA kit (ribodepletion workflow) and sequenced on an Illumina NovaSeq 6000 to generate 2 × 150 bp paired-end reads, targeting ∼50 million paired-end reads per sample. CPTAC-ccRCC isoform level FPKM and accompanying clinical metadata were obtained from the GDC database.

### 2.3. RNA-seq Preprocessing and Quantification

All raw sequencing reads were processed using a standardized Nextflow pipeline(v24.10.5)^28^. Adapter sequences and low-quality bases were removed using TrimGalore!, and read quality metrics were summarized using FastQC (RRID:SCR_014583) and MultiQC (RRID:SCR_014982). Reads were aligned to the human reference genome GRCh38.p13 using STAR (RRID:SCR_004463)(v2.7)^29^. Gene-level counts were quantified from aligned BAM files using featureCounts. Transcript-level quantification was performed using Salmon^30^ (v1.5) in mapping-based mode against the Ensembl v113 (RRID:SCR_002344) transcriptome to generate transcript-level estimated counts and TPM values. For downstream analyses requiring normalized expression, TPM values were log2-transformed as log2(TPM + 1). To identify potential sample contamination, Kraken2 and Bracken were used to screen reads for non-human content. Samples exceeding 1% bacterial, viral, or fungal reads were excluded prior to downstream statistical analyses.

### 2.4. DNA Methylation-Based Quality Control and Cell-Type Deconvolution

For DCC samples with matched Illumina HumanMethylationEPIC v1 profiles, raw IDAT files were preprocessed using the ENmix^31^ pipeline. Cross-reactive probes, SNP-associated probes, sex chromosome probes, and non-CpG probes were masked prior to downstream analysis. Probes with low-quality signal in more than 20% of samples within a given tissue type, defined as pOOBHA > 0.05, were removed. Histologic classification was verified using the classifier, HITAIC (RRID:SCR_028181), to exclude non-ccRCC samples. The tumor microenvironment composition was then inferred from methylation data using the HiTIMED (RRID:SCR_028180)^32^ deconvolution method, generating sample-level estimates of tumor, immune, and stromal fractions. These estimates were incorporated as covariates in adjusted differential expression and differential transcript usage models to account for variation in cellular composition.

### 2.5. Differential Gene Expression Analysis

Differential gene expression (DGE) analysis was conducted in R (v4.5) using the limma-voom^33^ method (RRID:SCR_010943), which models RNA-seq count data using precision-weighted linear modeling. The design matrix incorporated clinical and TME covariates: age, sex, and immune and angiogenic cell proportions. To account for potential confounding due to repeated measures or subject-level variability, a random effect for subject was included using the *duplicateCorrelation* function within the limma framework. Linear models were fit for each gene using the *lmFit* function, followed by empirical Bayes moderation of standard errors with *eBayes* to improve statistical stability. Genes were considered significantly differentially expressed if they met a Benjamini–Hochberg false discovery rate (FDR) adjusted p-value < 0.05 and an absolute log2 fold change (|logFC|) > 1.5. To contextualize DCC tumors relative to previously described IMmotion150/151 molecular programs, we also performed Non-negative matrix Factorization (NMF)-based clustering using gene signatures including angiogenesis, T-effector signaling, stromal activation, myeloid inflammation, complement cascade, fatty acid oxidation and metabolism, cell cycle, and omega oxidation^10^. Signature scores were computed across tumor samples, and consensus stability metrics were used to select optimal k clusters. Median z-scored program activity was then compared across clusters.

### 2.6. Batch Correction and Consensus Clustering of Transcript-Level Expression

Transcript-level expression matrices (log2(TPM+1)) were batch-corrected using the removeBatchEffect function from limma, with batch specified as the primary technical factor and additional covariates included (age, sex, immune and angiogenic proportions). For clustering, the top 1000 variable transcripts were selected based on variance across tumor samples (selection threshold specified in the analysis scripts), and consensus clustering was performed using the ConsensusClusterPlus^34^ framework. Cluster number (k) was determined based on the cumulative distribution function (CDF) behavior. Cluster assignments were used for downstream DTU comparisons and association testing with clinical and microenvironment features.

### 2.7. Differential Transcript Usage Analysis

Transcript-level estimated counts generated by Salmon (approximately 387,944 transcripts) were first filtered to remove low-count features. For the tumor versus normal comparison, transcripts were retained if they had at least 10 counts in 33 samples, corresponding to the size of the smaller comparison group. Filtering was performed using min_samps_feature_expr = 33, min_gene_expr = 10, and min_feature_expr = 10. After filtering, 52,154 transcripts remained. For testing differences in relative isoform usage between conditions (tumor versus normal), differential transcript usage (DTU) was evaluated using a two-stage framework. This staged approach combined Dirichlet-multinomial modeling for initial DTU detection implemented in DRIMSeq^35^, with limmavoom modeling for downstream covariate-adjusted effect estimation. This approach aimed to overcome convergence limitations of the DRIMSeq framework when fitting covariate-rich multivariable models. The first stage used Dirichlet–multinomial modeling implemented in DRIMSeq to test for differential transcript usage between groups while accounting for batch effects. Specifically, DRIMSeq’s *dmFilter* function was applied to remove genes with only one transcript (“lonely transcripts”), resulting in 45,564 transcripts entering model fitting.

Following DTU testing, gene-level screening p-values and transcript-level confirmation p-values were integrated using StageR (v1.20) to control the overall gene-level false discovery rate. Transcripts meeting StageR-adjusted significance thresholds (FDR < 0.05 unless otherwise specified) were retained for downstream analysis in the second stage.

The second stage implemented covariate-adjusted modeling using the weighted least squares linear modeling framework *limma-voom*. To obtain covariate-adjusted effect estimates and evaluate the influence of clinical and tumor microenvironment covariates, the significant transcripts identified in the first stage were modeled with adjustment for age, sex, AUA risk group, and proportions of immune and angiogenic tumor microenvironment components. The linear model used for effect estimation was:

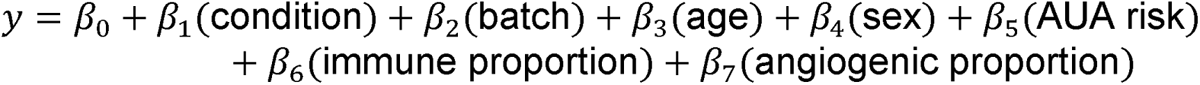

An analogous DTU workflow was applied to comparisons between the two tumor clusters defined by consensus clustering. For the within-tumor comparison, transcripts were retained if they had at least 10 counts in 35 samples, corresponding to the size of the smaller comparison group. Filtering was performed using min_samps_feature_expr = 35, min_gene_expr = 10, and min_feature_expr = 10. In this analysis, 59,898 transcripts remained after initial filtering, and 52,048 transcripts were retained following removal of lonely transcripts prior to Dirichlet–multinomial modeling. For the second stage of the cluster comparison, the covariate-adjusted linear model used for effect estimation was:

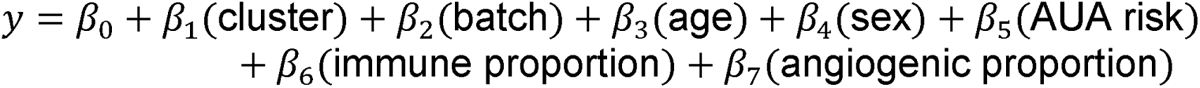

Final differentially used transcripts were defined as transcripts exhibiting statistically significant and directionally concordant changes across both the DRIMSeq/StageR DTU modeling and second-stage covariate-adjusted limma-voom modeling (Benjamini-Hochberg FDR < 0.05 and |logFC| > 1.5).

### 2.8. Functional Annotation and Gene Set Enrichment Analyses

Functional enrichment analyses were conducted using clusterProfiler (v4.6) (RRID:SCR_016884) in R. For DEG results, Gene Set Enrichment Analysis (GSEA) was conducted using *gseGO* with signed gene-level statistics, focusing on Gene Ontology Biological Process terms and the c5 hallmark category and hallmark terms for epithelial-to-mesenchymal transition, hypoxia, and fatty acid metabolism.

Multiple testing correction was applied using the Benjamini–Hochberg procedure, with significance defined at FDR < 0.01 unless otherwise stated. For DTU analyses, transcript-level results were first mapped to their corresponding gene identifiers prior to enrichment analysis. Significant transcripts were stratified according to transcript biotype: protein-coding transcripts were categorized as functional, and transcripts annotated as non-sense-mediated decay, retained intron, or protein-coding CDS not defined were categorized as non-functional. Within each biotype category, transcripts were further stratified by differential usage direction based on covariate-adjusted log fold change estimates, and Gene Ontology over-representation analysis was performed separately for each sub-group. To prevent inflation arising from multiple transcripts mapping to the same gene, a maximum-weighted selection approach was implemented. For each gene within a given biotype category, the transcript with the largest absolute effect size (|LogFC|) was retained while preserving the direction of effect. This yielded a single representative statistic per gene per category. The resulting gene lists were used as input for gene set enrichment analysis (GSEA), which was performed independently for functional and non-functional transcript groups. Gene Ontology (GO) over-representation analysis was also performed per biotype group using *enrichGO*. Protein domain information were annotated using pfam (http://pfam.xfam.org/).

### 2.9. Prognostic Model Construction and Internal Validation

To construct a transcript-based prognostic model, log2-normalized expression of DTU-derived candidate transcripts (defined by the prespecified DTU significance threshold) was evaluated using Cox proportional hazards regression with LASSO regularization implemented in *glmnet*^38^. Internal validation was performed using bootstrap resampling with a 70:30 train–test split, repeated over 200 times. In each iteration, a Cox-LASSO model was trained on the training set and evaluated on the held-out test set. In parallel, a clinical-only Cox-LASSO model incorporating age, sex, and AUA risk category was fit to provide a baseline comparator. Feature stability was assessed by quantifying selection frequency across bootstrap iterations, and predictors selected more than 10 times were retained as stable features. Transcript-based and clinical-based risk scores were generated from the glmnet-derived linear predictors, and a joint risk score was computed by summing these components. Final model coefficients were refit on the full cohort using the retained predictors. The resulting composite risk score, integrating transcript usage with clinical variables, was used to stratify patients into high- and low-risk groups based on the median score. Survival differences were evaluated using Kaplan–Meier analysis, and associations with overall survival were assessed using multivariable Cox proportional hazards models adjusting for age, sex, and AUA risk category.

### 2.10 Statistical Analysis

Associations between cluster membership and categorical clinical features (e.g., AUA risk group) were evaluated using chisquare tests. Linear models were adjusted for age, sex and tumor micro environment composition. Differences in continuous variables (e.g., immune and angiogenic proportions) were assessed using Wilcoxon rank-sum tests (implemented from the R package, stats v4.5.3). Prognostic model performance was summarized using Harrell’s C-index, and nested model comparisons were performed using likelihood ratio tests where appropriate. All analyses were performed in R (v4.5). Continuous variables are reported as median (interquartile range), and categorical variables are reported as counts (percentages). Unless otherwise noted, a two-sided p < 0.05 was considered statistically significant. Code and processed data are available at https://github.com/SalasLab/DTU-Analysis-in-ccRCC-Reveals-Isoform-Level-Remodeling.

## 3. Results

### 3.1. Study Population

The DCC cohort comprised 127 ccRCC tumors with available clinical data for 109 patients, alongside 33 adjacent normal tissues. Patients were predominantly male (64.2%) with a mean age of 60.9 years. Most tumors were early stage, with stage I being the most common (47.7%), followed by stage III (22.9%), stage IV (16.5%), and stage II (12.8%). Tumor grade was largely intermediate to high, with Grade 2 (40.4%) most frequent, followed by Grade 3 (28.4%) and Grade 4 (20.2%), while Grade 1 tumors were uncommon (4.6%). Consistent with this, most patients (67.0%) were classified as intermediate- or high-risk by AUA criteria. At follow-up, 40.4% of patients experienced an event, with a median overall survival of 1.72 years. (**Table 1)**

**Table 1.** Clinical and pathologic characteristics of the Dartmouth Cancer Center (DCC) ccRCC cohort. (Adjacent normal, n=33; 28 complete cases)

| Groups | Overall | Censored | Event | p |
| --- | --- | --- | --- | --- |
| Number of tumors =127; 109 complete cases) | 109 | 65 | 44 |  |
| **Age, mean (SD)** | 60.94 (11.23) | 60.48 (10.29) | 61.61 (12.49) | 0.608 |
| **sex (%)** |  |  |  | 0.003 |
| Male | 70 (64.2) | 34 (52.3) | 36 (81.8) |  |
| Female | 39 (35.8) | 31 (47.7) | 8 (18.2) |  |
| **Stage (%)** |  |  |  | <0.001 |
| I | 52 (47.7) | 39 (60.0) | 13 (29.5) |  |
| II | 14 (12.8) | 10 (15.4) | 4 (9.1) |  |
| III | 25 (22.9) | 12 (18.5) | 13 (29.5) |  |
| IV | 18 (16.5) | 4 (6.2) | 14 (31.8) |  |
| **Grade (%)** |  |  |  | **0.021** |
| I | 5 (4.6) | 5 (7.7) | 0 (0.0) |  |
| II | 44 (40.4) | 31 (47.7) | 13 (29.5) |  |
| III | 31 (28.4) | 18 (27.7) | 13 (29.5) |  |
| IV | 22 (20.2) | 9 (13.8) | 13 (29.5) |  |
| **AUA Risk, binary (%)** |  |  |  | **0.037** |
| Intermediate/high | 73 (67.0) | 38 (58.5) | 35 (79.5) |  |
| Low | 36 (33.0) | 27 (41.5) | 9 (20.5) |  |

### 3.2. Differential Gene Expression Analysis with Adjustment for Cell-Type Confounders reveals characteristic differential immune and metabolic patterns in ccRCC

Differential gene expression analysis adjusting for HiTIMED-derived immune and angiogenic cell-type proportions identified 2,176 significantly differentially expressed genes (1,031 upregulated and 1,145 downregulated) between ccRCC tumors and adjacent normal tissues (Figure 1A). Among the most strongly upregulated genes were *CA9*, a hypoxia-inducible carbonic anhydrase; *FABP7,* involved in lipid binding, and *SLC6A3*, the dopamine transporter^40–42^. These were observed alongside established markers of tumor-associated signaling and epithelial–mesenchymal transition.

**Figure 1.**
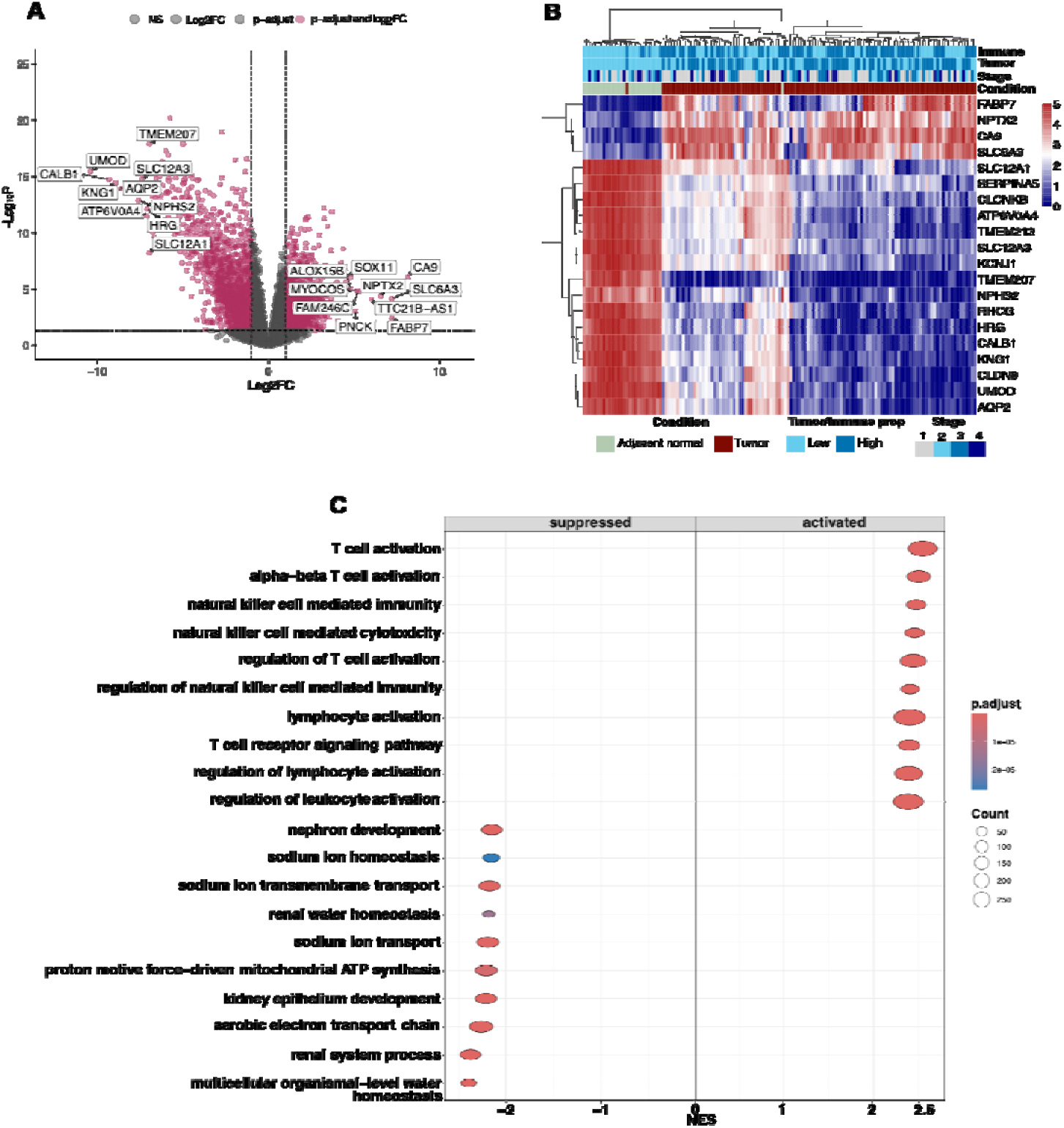
Differential Gene Expression Analysis with Adjustment for Cell-Type Confounders reveals characteristic differential immune and metabolic patterns in ccRCC. **(A)** Volcano plot showing differential gene expression between ccRCC tumors and adjacent normal kidney tissues after adjustment for immune and angiogenic cell-type proportions. The x-axis represents log_2_ fold change, and the y-axis represents −log_10_ adjusted p-value. Significantly upregulated and downregulated genes are highlighted. **(B)** Heatmap of top 20 differentially expressed genes across tumor and normal samples. Rows represent genes and columns represent samples. Expression values are scaled per gene. **(C)** GSEA analysis of differentially expressed genes. Dot plots display enriched biological processes separately for upregulated (activated) and downregulated (suppressed) genes. The x-axis represents gene ratio, dot size corresponds to gene count, and color indicates adjusted p-value.

In contrast, the most prominently downregulated genes included those involved in renal function, such as *UMOD*, *CALB1*, *KNG1*, *AQP2*, *NPHS2,* and *SLC12A3*^43,44^. (Figure 1B). GSEA revealed dysregulation of characteristic immune, renal, and metabolic function pathways (Figure 1C). Genes upregulated in tumors were enriched for immune-related biological processes, including T cell activation, natural killer cell-mediated immunity, cytokine production, and regulation of leukocyte activation. In contrast, downregulated genes were enriched for pathways related to renal structure and function and mitochondrial function, including nephron development, mitochondrial ATP synthesis, and aerobic electron transport chain. Hallmark pathway analysis demonstrated enrichment of epithelial–mesenchymal transition and hypoxia signatures among upregulated genes, whereas metabolic pathways, including fatty acid metabolism, were enriched among downregulated genes (Supplementary Figure 1A-B). Finally, NMF-based program scoring resolved four stable transcriptomic clusters based on consensus stability metrics. Although this four-cluster solution did not fully recapitulate the granularity of the IMmotion subtypes, Cluster 4 showed strong enrichment for angiogenesis and FAO/AMPK-related metabolic programs, consistent with the angiogenic subtype described in the IMmotion analyses^10^. The remaining clusters showed relative enrichment of immune-inflammatory, stromal, or proliferative features. (Supplementary Figure 1C-D).

### 3.3. Differential Transcript Usage reveals enrichment of pathways related to renal function, immune response, and altered metabolism among splice variants

StageR filtering of DRIMSeq output identified 9,623 genes significant at the first stage. Part 2 of the DTU analysis identified 1,170 transcripts with significant differences in usage between ccRCC tumors and adjacent normal tissues (Figure 2A, Supplementary Figure 2A), mapping to 881 genes. Of these, 532 transcripts showed increased usage in tumors, whereas 638 showed decreased usage. 182 genes were common between genes with Differential Transcript Usage (gDTUs) and DEGs, including the top 10 DEGs (Supplementary Figure 1E)

**Figure 2.**
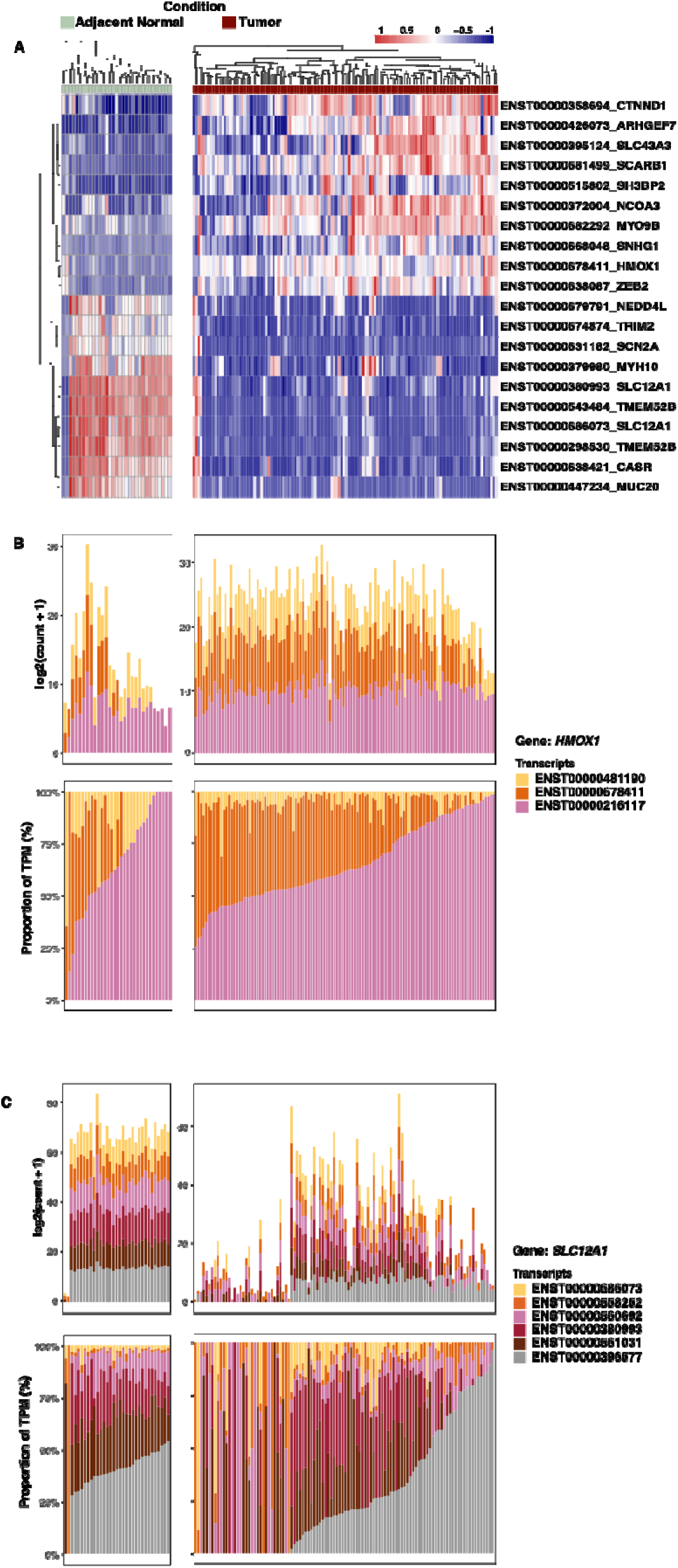
D Differential Transcript Usage reveals enrichment of pathways related to renal function, immune response and altered metabolism among splice variants. **(A)** Heatmap of top 10 transcripts with increased/decreased usage (|logFC >1.5|, FDR < 0.05). **(B-C)** Stacked bar plots showing isoform usage proportions for top two genes with DTU (B) top gene with increased (C) top gene with decreased usage. Each bar represents a sample, the y-axis represents log2 normalized counts and transcript proportion, and the x-axis represents samples in either condition.

The transcripts with the greatest increases in usage included transcripts for *ENST00000678411* (*HMOX1*), a heme oxygenase involved in the oxidative stress response; *ENST00000681499* (*SCARB1*), a scavenger receptor mediating cholesterol uptake; and *ENST00000358694* (*CTNND1*), a regulator of cell adhesion and Wnt signaling. In contrast, the most pronounced decreases were observed for *ENST00000686073* (*SLC12A1*), a sodium-potassium-chloride cotransporter critical for renal ion transport, and *ENST00000298530* (*TMEM52B*), associated with epithelial structure and membrane organization^48,49^ (Figure 2B–C). For these loci, transcript usage direction was consistent with gene-level differential expression (Supplementary Figure 2B–D).

Given the strong decrease observed for *SLC12A1*, we evaluated the potential impact of low-count samples. Two normal samples, RCC447N and RCC574N, exhibited particularly low counts, and several additional samples showed relatively low read depth despite passing quality control. Sensitivity analyses excluding eight such samples did not materially alter the results (Supplementary Table 1).

Comparison of transcript-level and gene-level results identified three transcripts across two genes exhibiting discordant regulation. For *FN1*, which is involved in extracellular matrix organization, overall gene expression increased in tumors, whereas *FN1:ENST00000357867* (SE) showed decreased usage, indicating isoform-specific repression within an otherwise upregulated gene^50^. Conversely, for *SLC8A1*, a solute carrier, gene-level expression decreased in tumors, while *SLC8A1:ENST00000705281* (AF) and *SLC8A1:ENST00000406785* (SE) both showed increased usage^50,51^ (Supplementary Table 1).

Weighted gene set enrichment analysis of StageR-significant transcripts from the first-stage model, comprising 5,234 functional and 3,254 non-functional isoforms, revealed distinct pathway patterns associated with transcript usage across both transcript classes. **(Figure 3A–C).** Among transcripts in the functional biotype, positively enriched Gene Ontology biological process terms included immune-related pathways such as B cell activation, B cell differentiation, and lymphocyte proliferation. In contrast, pathways related to ion transport and transmembrane exchange were negatively enriched. In contrast, non-functional biotype transcripts exhibited positive enrichment of protein metabolic and post-transcriptional regulatory processes, whereas lipid metabolic pathways, including fatty acid and lipid oxidation, were negatively enriched. Similarly, overrepresentation analysis largely recapitulated similar pathway distribution between biotypes. (**Supplementary Figure 3)**.

**Figure 3.**
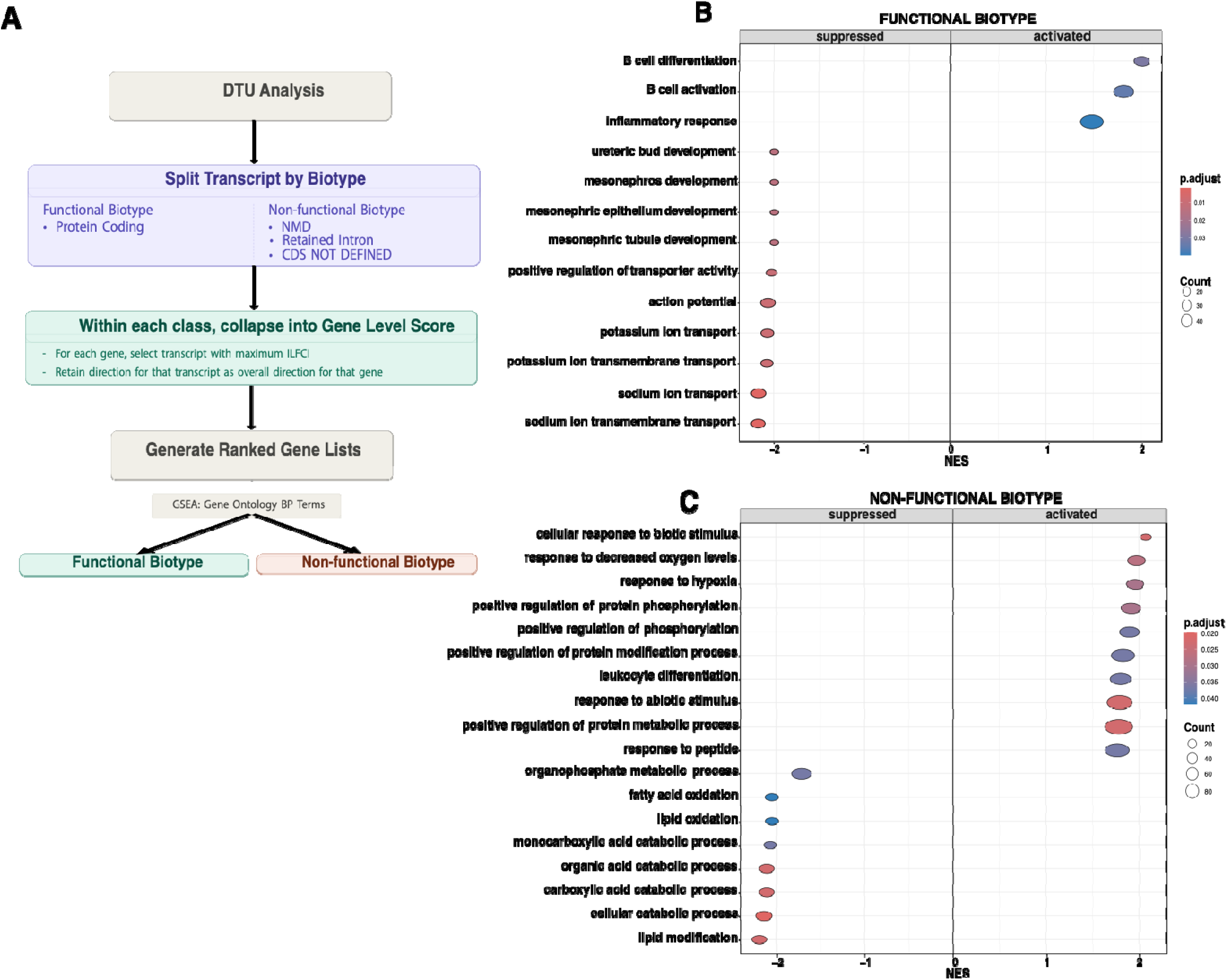
Transcript-level reprogramming in ccRCC reveals coordinated functional and non-functional pathway remodeling. (A) Schematic overview of the transcript-level enrichment workflow. Differential transcript usage (DTU) results were stratified by transcript biotype into functional (protein-coding) and non-functional (e.g., nonsense-mediated decay, retained intron, CDS not defined) classes. Within each class, transcripts were collapsed to gene-level scores by selecting the transcript with the largest absolute log fold change (|logFC|), retaining directionality. Ranked gene lists were then generated and used for gene set enrichment analysis (GSEA). (B-C) GSEA on Gene Ontology (GO) Biological processes (BP) Terms for functional and non-functional (C)transcripts. Dot plots show significantly enriched biological processes among transcripts with increased (activated) or decreased (suppressed) usage in tumors relative to normal tissue. Dot size represents gene set size and color indicates adjusted p-value.

### 3.4. Consensus Clustering of Isoform Expression Identifies Two Tumor Subgroups with differential patterns of transcript usage

Consensus clustering based on transcript-level expression profiles identified two tumor clusters (k = 2), as supported by the CDF and delta area plots (Figure 4A, Supplementary Figure 4A). Cluster 1 was enriched for tumors classified as high risk by AUA binary stratification and showed higher angiogenic cell-type infiltration scores than Cluster 2 (Supplementary Figure 4B, Supplementary Table 2). Differences in immune-related features were also observed between clusters, although these did not reach statistical significance. Kaplan-Meier survival analysis demonstrated a separation in overall survival between the two transcript-defined clusters (log-rank P = 0.0076; Figure 4B). By comparison, stratification by AUA binary risk alone also showed significant survival differences, but with less pronounced separation in the same cohort (Figure 4C).

**Figure 4.**
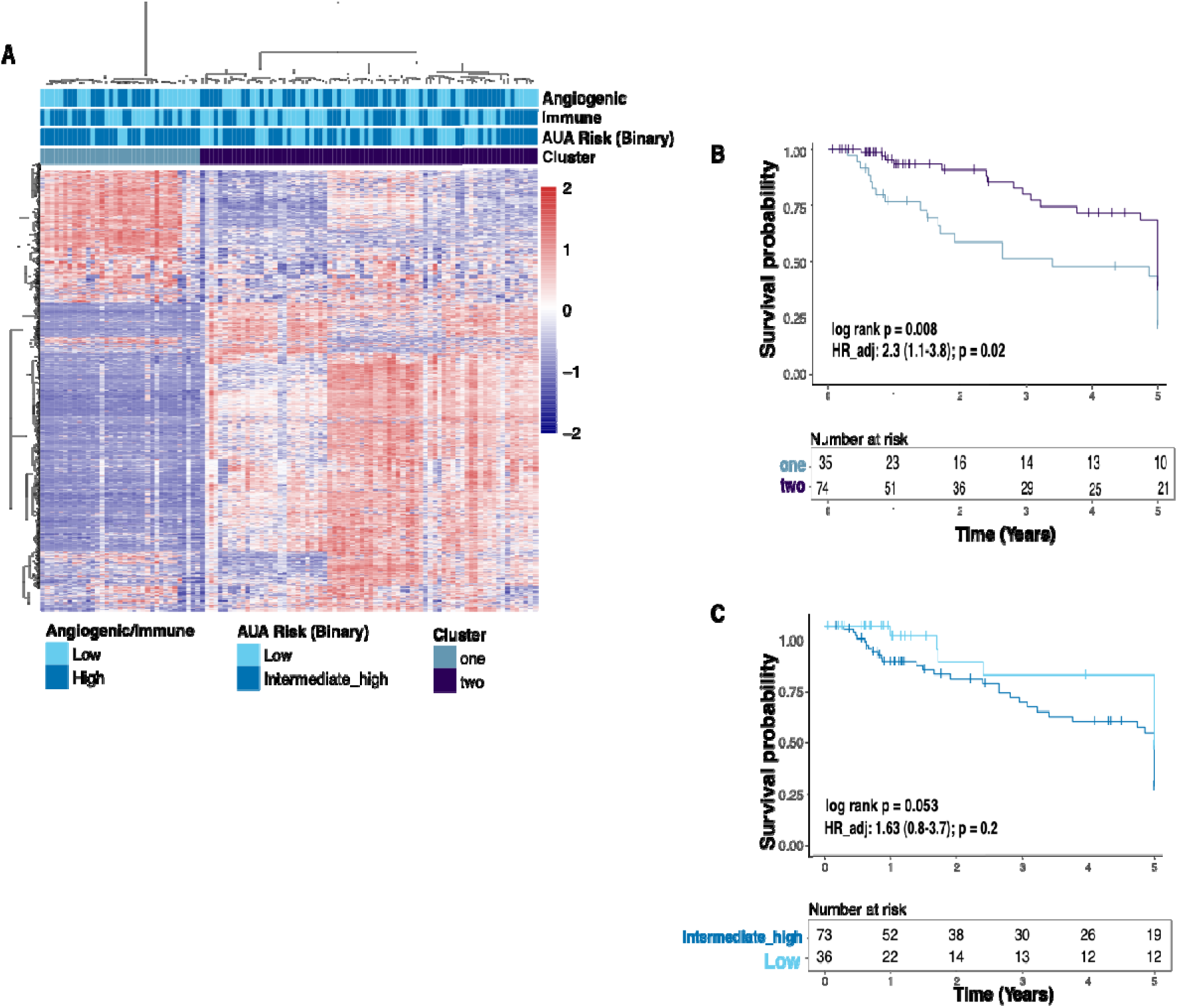
Consensus Clustering of Isoform Expression Identifies Two Tumor Subgroups with differential patterns in transcript usage. (A) Heatmap of transcript-level expression profiles showing two clusters identified by consensus clustering, with annotation tracks for angiogenic and immune scores, and AUA binary risk category. (B) Kaplan–Meier overall survival curves comparing patients in cluster 1 versus cluster 2. Survival estimates were derived from Cox proportional hazards models adjusted for age, sex, and AUA risk category (log-rank p-value shown). (C) Kaplan–Meier overall survival curves stratified by AUA binary risk category (low vs intermediate/high), adjusted for age, sex, and stage (log-rank p-value shown).

Differential transcript usage analysis comparing the two tumor clusters identified 19,595 transcripts with significant usage differences, including 10,470 with increased usage and 9,123 with decreased usage, corresponding to 7,940 genes (Figure 5A, Supplementary Table 3). Among transcripts with increased usage, top-ranking features included transcripts from genes involved in cytoskeletal organization, such as *ACTB: ENST00000676319 (AF),* and immune-related function, including *B2M: ENST00000557901 (AF; NMD*)^52,53^. Importantly, compared to their canonical isoforms, *ACTB: ENST00000676319* lacks the ASKHA_NBD_actin domain, while *B2M:ENST00000557901* lacks the immunoglobulin C1-set domain (Supplementary Table 4). Transcripts with decreased usage included those derived from genes associated with extracellular matrix organization and cell motility, such as *FN1: ENST00000356005 (A3)* and *APLP2*: ENST00000528499 (A5/AF), as well as iron metabolism, including *FTH1: ENST00000530019* (AL). Additional domain-level differences were observed among FN1 isoforms; for example, *FN1:ENST00000456923* lacks both N- and C-terminal signal peptide domains while retaining a fibronectin type III repeat signature (Supplementary Table 4). Several recurrent patterns were evident among the top differentially used transcripts. *FN1* contributed two non-canonical protein-coding transcripts among the top five increased and decreased usage features, highlighting marked isoform redistribution within the same gene. Multiple protein-coding *ACTB* transcripts were also represented among the most strongly altered features, including *ACTB: ENST00000473257*(A3, SE) in the top three and *ACTB: ENST00000464611* (AL, SE) within the top 15. Similarly, protein-coding *FTH1* contributed more than one transcript in both directions, including *FTH1: ENST00000526640* (A3) (Increased Usage) and *FTH1:ENST00000530019* (SE) (Decreased Usage). *EGFR: ENST00000450046* (SE) was also among the top five transcripts with decreased usage.

**Figure 5.**
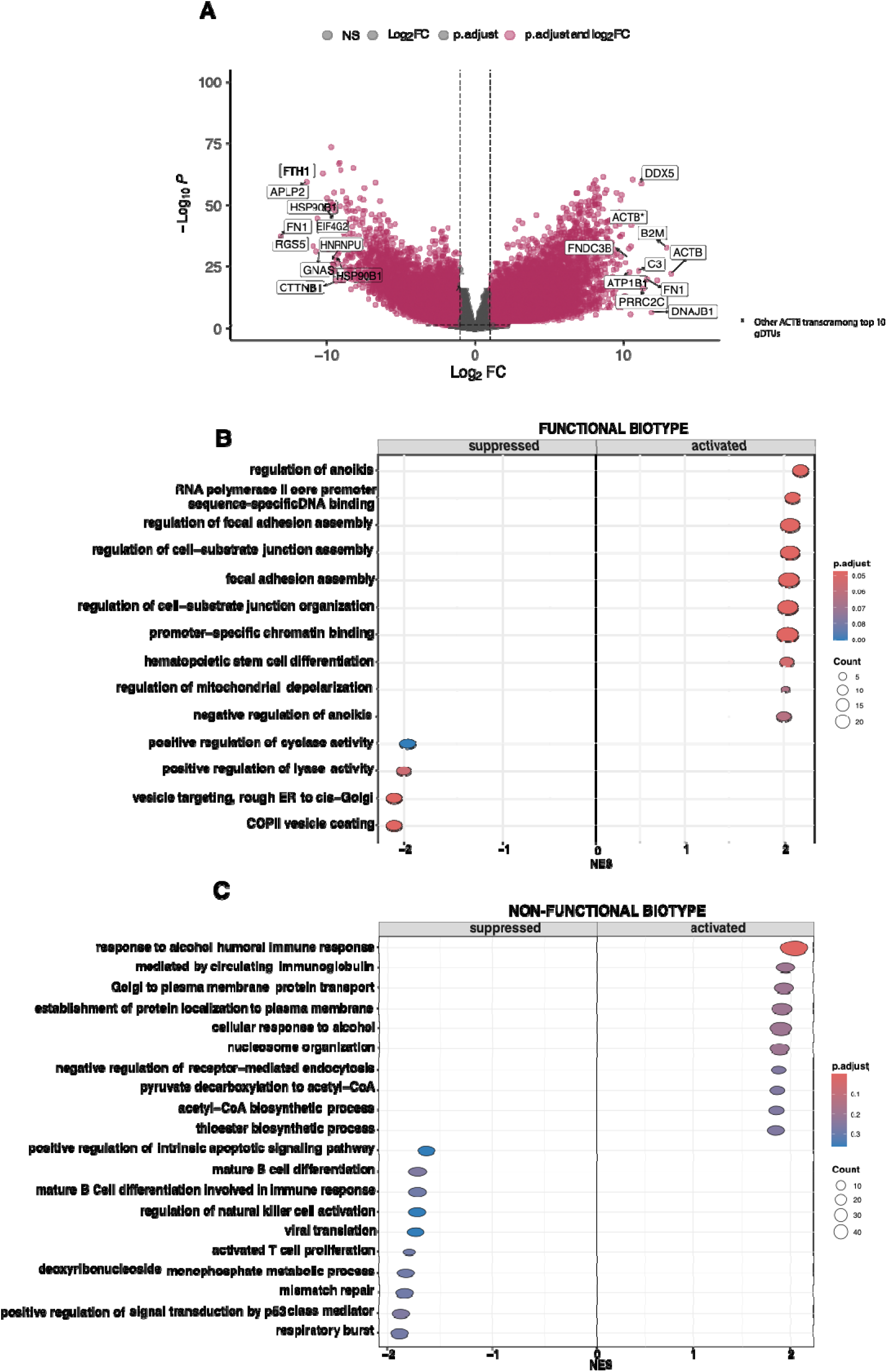
Tumor cluster–specific transcript usage highlights transcriptional activation and metabolic suppression programs. **(A)** Volcano plot of differential transcript usage showing log_□ fold change versus −log□□ adjusted P value. Top 10 Transcripts for each direction are labeled. **(B–C)** Weighted gene set enrichment analysis results stratified by transcript biotype. (B) Functional (protein-coding) transcripts. **(C)** Non-functional transcripts.

Weighted gene set enrichment analysis of StageR-significant transcripts from the first stage model, comprising 20,649 functional and 12,598 non-functional isoforms, revealed distinct pathway architectures associated with transcript usage across both transcript classes (Figure 5B-C). For functional transcripts, increased usage was enriched for processes related to transcriptional regulation, including RNA polymerase II promoter-specific DNA binding, chromatin binding, miRNA transcription and regulation, as well as vesicle trafficking and cell-substrate junction organization. In contrast, decreased usage of functional transcripts was enriched for pathways involving regulation of anoikis, cyclase activity, and vesicle transport. For non-functional transcripts, increased usage was enriched for regulatory and signaling processes, including positive regulation of cyclase activity, response to prostaglandin E, metabolic compound salvage, and pathways related to methylation and vesicle trafficking. Conversely, decreased usage of non-functional transcripts was enriched in metabolic and signaling pathways, including membrane lipid biosynthesis, lipid metabolic processes, *cGAS-STING* signaling, and cellular responses to external stimuli.

### 3.5. Prognostic Risk Stratification Derived from Differentially Used Transcripts

Our modeling framework identified twelve transcripts that were consistently retained across iterations. **(Figure 6A).** These features spanned multiple transcript biotypes, including protein-coding, retained intron, nonsense-mediated decay, long non-coding RNA, and TEC-classified isoforms, highlighting the contribution of both canonical and non-canonical transcripts. **(Table 2).**

**Figure 6.**
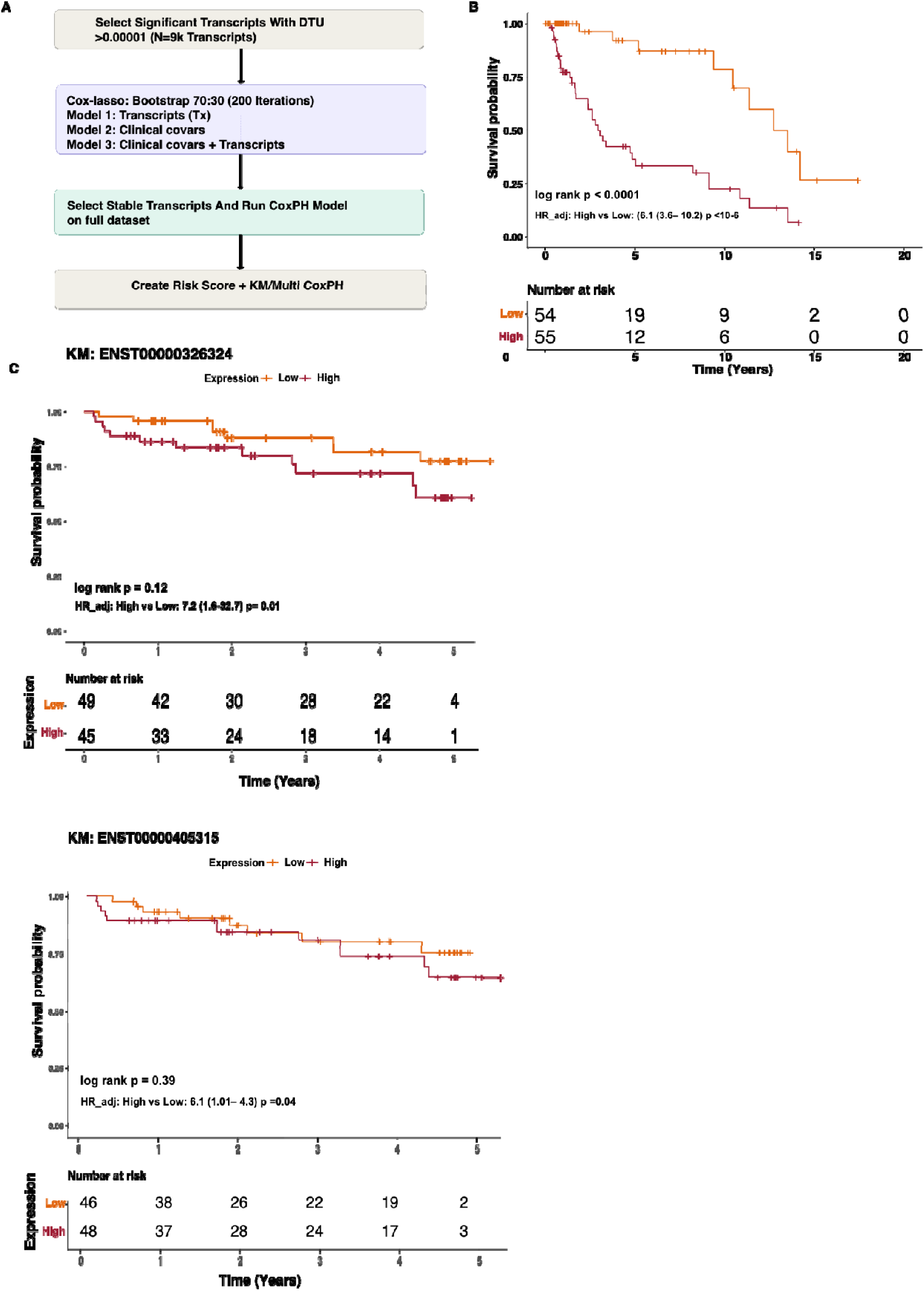
Prognostic Risk Stratification Derived from Differentially Used Transcripts. **A)** Workflow for prognostic model construction. Transcripts significant in DTU analysis between tumor clusters (StageR-adjusted FDR < 0.05) were entered into Cox proportional hazards modeling with LASSO regularization using 70:30 bootstrap resampling (200 iterations). Transcript-only and clinical-only models were compared, and a composite risk score was generated from stable transcript and clinical features. **(B)** Kaplan–Meier overall survival curves stratified by composite risk score (median split). Log-rank test p value is shown and multivariable Cox proportional hazards model adjusted for age, sex, and AUA risk category are shown. Hazard ratio (HR) with 95% confidence interval is reported. Table 2. Stable transcripts retained in bootstrap Cox–LASSO modeling, including transcript ID, external gene name, transcript biotype, annotated function, and canonical status. **(C)** Validation of two of the twelve prognostic DTUs in the CPTAC-ccRCC cohort.

**Table 2.**
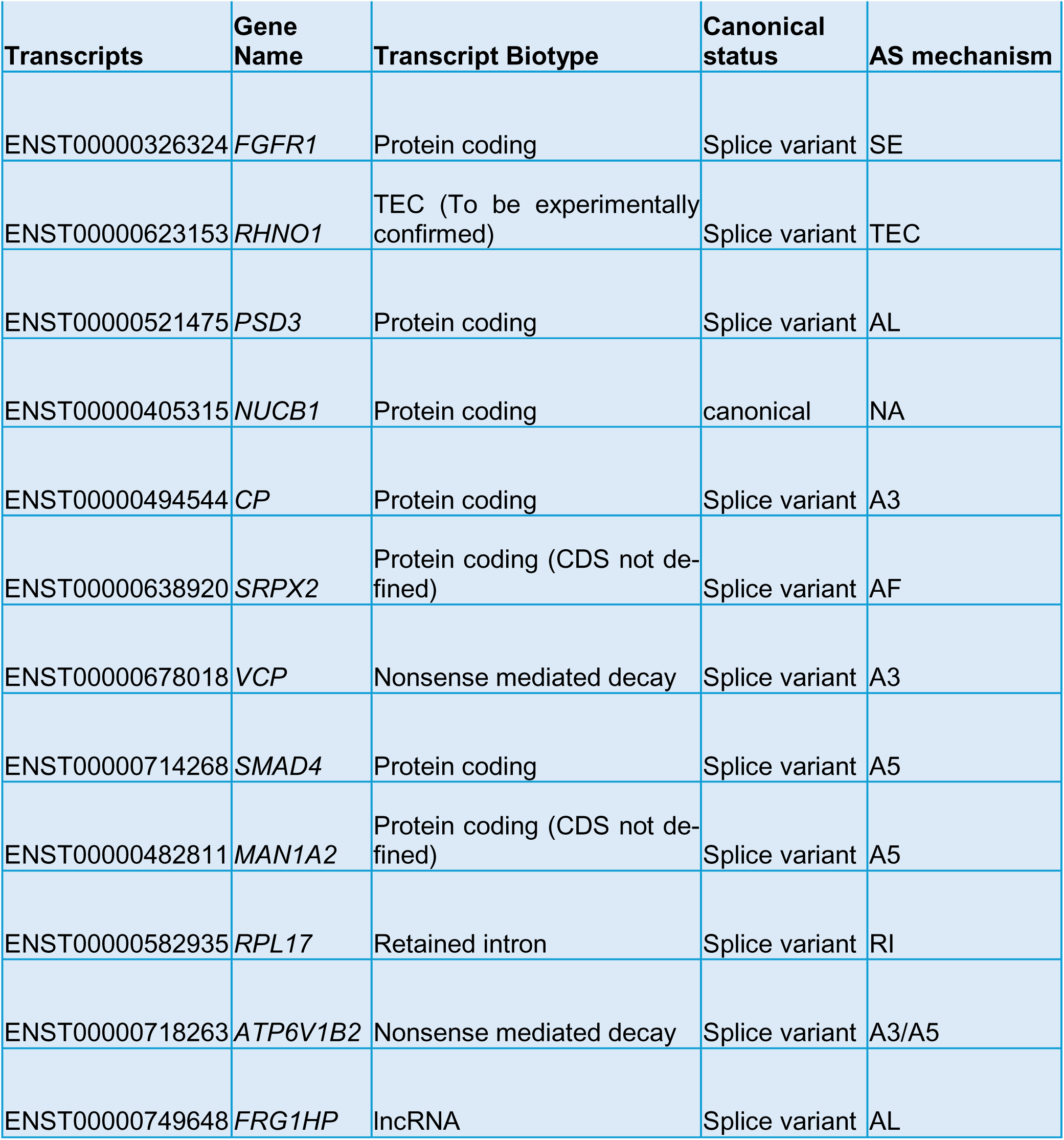
Prognostic DTU-Derived Transcripts with Associated Gene, Biotype, and Alternative Splicing Mechanism in ccRCC.

Functionally, the retained transcripts represented diverse biological processes relevant to ccRCC. These included angiogenesis (*FGFR1*, *ENST00000326324*; A3), DNA damage response (*RHNO1*, *ENST00000623153*;A3), vesicle trafficking (*PSD3*, *ENST00000521475*; A5; *CP*, *ENST00000494544*; A3), extracellular matrix remodeling (*SRPX2*, *ENST00000638920*; ES), protein homeostasis (*VCP*, *ENST00000678018*; A5), TGF-β signaling (*SMAD4*, *ENST00000714268*; A5), glycoprotein processing (*MAN1A2*, *ENST00000482811*; A5), translation (*RPL17*, *ENST00000582935*; RI), and vesicular acidification (*ATP6V1B2*, *ENST00000718263*; A3)^56–63^. In addition, a long non-coding RNA (*FRG1HP*, *ENST00000749648*; AL) and a calcium-associated protein-coding isoform (*NUCB1*, *ENST00000405315*; ES) were identified among the prognostic features^64^. Integration of transcript-derived features with clinical variables (age, sex, and AUA risk category) improved risk stratification compared to clinical variables alone (Supplementary Figure 5). Kaplan–Meier analysis demonstrated significantly worse overall survival in the high-risk group defined by the composite score (log-rank p < 0.0001) **(Figure 6B).** External validation in the CPTAC ccRCC cohort was limited by transcript overlap, with only two of the twelve transcripts available. However, both *ENST00000326324* (*FGFR1*) and *ENST00000405315* (*NUCB1*), including a non-canonical protein-coding isoform, remained independently associated with survival after adjustment for clinical covariates **(Figure 6C).**

## 4. Discussion

In this study, we integrated gene-level expression, differential transcript usage, tumor microenvironment adjustment, transcript-based clustering, and prognostic modeling to comprehensively characterize the ccRCC transcriptomic landscape. Our findings show that transcript usage captures tumor biology that is not fully reflected in aggregate gene expression alone. While adjusted gene-level results recapitulated known ccRCC hallmarks, isoform-level analysis revealed extensive transcript remodeling.

For the tumor vs adjacent normal comparison, most transcripts with significant DTU mirrored gene-level directionality; however, three transcripts exhibited discordant patterns relative to overall gene expression, suggesting that transcript remodeling in ccRCC is largely coordinated with gene-level changes but can reveal additional regulatory complexity. For *FN1*, increased gene expression alongside decreased usage of *FN1:ENST00000357867* (SE) suggests isoform-specific remodeling of extracellular matrix composition rather than uniform upregulation, consistent with functional diversification of fibronectin isoforms in tumor invasion^50,65^. Conversely, for *SLC8A1*, reduced gene-level expression with increased usage of *SLC8A1:ENST00000705281* (AF) and *SLC8A1:ENST00000406785* (SE) may indicate selective preservation of specific isoforms. This pattern may reflect preservation of calcium signaling capacity under global repression, or alternatively, the emergence of isoforms with distinct regulatory roles in calcium homeostasis that could also contribute to tumor-promoting signaling^51^.

Several isoform switches further support transcript-level remodeling of canonical ccRCC pathways. The pronounced loss of *SLC12A1* and related solute carrier transcripts suggests that nephron identity loss extends beyond reduced gene expression to include altered isoform usage within renal transport genes^66^. Similarly, decreased usage of a *TMEM52B* alternative last-exon isoform and increased usage of exon-skipping *HMOX1* isoforms point to remodeling of epithelial integrity and oxidative stress adaptation^45,49^. Increased usage of a retained-intron *SCARB1* isoform further suggests post-transcriptional regulation of lipid metabolism^46^. Despite *SCARB1’s* established tumor-promoting role in lipid metabolic adaptation, this non-productive isoform may fine-tune functional *SCARB1* output, enabling tighter control of lipid handling rather than simple pathway activation. The biological relevance of these findings is strengthened by consistency with prior studies. Among the DTUs were splice variants for genes previously reported as tumor-associated, including *SLC15A4* and *SYNPO*, although these variants did not show comparable prognostic significance in our cohort^20^. The detection of an *EGFR* splice variant in our cohort is also consistent with reports of aberrant *EGFR* splicing that have been linked to aggressive disease and therapeutic resistance^21^.

Stratification by transcript biotype revealed distinct but complementary regulatory patterns. Functional, protein-coding transcripts primarily reflect activation of immune-related programs, consistent with the inflamed tumor microenvironment, whereas non-functional isoforms appear to capture suppression of metabolic pathways, including fatty acid and lipid oxidation. One potential explanation is that ccRCC leverages non-productive transcripts, such as NMD and retained intron isoforms, to attenuate post-transcriptional metabolic gene output, reinforcing the well-described metabolic shutdown at the gene level. In parallel, selective enrichment of functional isoforms within immune pathways may enhance effective protein production required for signaling and immune interaction. Together, this split suggests that alternative splicing enables coordinated regulation of tumor biology by simultaneously amplifying immune functions through productive isoforms while constraining metabolic processes via non-functional transcript usage. Given the close dependence of immune-cell activity on metabolic fitness, this discordance may reflect a state in which immune signaling appears transcriptionally activated, yet occurs in the context of impaired or dysregulated metabolic support^67,68^. More broadly, this dynamic reinforces the established coupling between metabolic rewiring and immune signaling in ccRCC, suggesting that isoform-level regulation coordinates these hallmark tumor processes^69^. Within-tumor analysis further highlighted the importance of isoform-level regulation. Transcript-based clustering identified two tumor subgroups with distinct survival outcomes and biological characteristics, outperforming AUA risk stratification alone. The transcript features distinguishing these cluster points to coordinated remodeling of core cellular systems.

Increased usage of *ACTB: ENST00000676319*, which lacks the ASKHA/ASKHA_NBD_actin domain relative to the canonical transcript, may indicate reduced preservation of core actin-associated functional architecture and could reflect remodeling of cytoskeletal dynamics rather than simple upregulation of *ACTB*^52^. Similarly, the enrichment of a *B2M* nonsense-mediated decay transcript suggests isoform-level modulation of antigen presentation, whereby increased usage of non-productive transcripts may reduce functional antigen-presenting capacity and contribute to immune evasion^53^. In contrast, decreased usage of *FN1:ENST00000456923*, which lacks the N- and C-terminal signal peptide domains while retaining a fibronectin type III repeat signature, suggests reduced expression of an isoform with altered secretion or extracellular localization, potentially shifting fibronectin composition toward variants that more effectively support matrix assembly, adhesion, and tumor invasion^50,70^. Additionally, decreased usage of *FN1, APLP2, FTH1,* and *EGFR* isoforms implicates extracellular matrix organization, iron metabolism, and receptor signaling as axes of divergence^21,50,54,55^. Altered *FTH1* transcript usage may reflect isoform-level regulation of iron sequestration and ferroptosis susceptibility within the metabolically rewired state of ccRCC. Given the emerging role of ferroptosis in linking lipid metabolism, oxidative stress adaptation, and immune signaling in renal cancer, these findings suggest that transcript remodeling may contribute to how tumors balance metabolic fitness with resistance to lipid peroxidation-driven cell death^71^. These findings indicate that alternative splicing contributes to intratumoral heterogeneity by fine-tuning the functional output of transcripts across key dysregulated pathways in ccRCC.

The prognostic modeling results further underscore that a clinically relevant signal resides at the transcript level. The retained transcripts spanned diverse biological processes central to ccRCC, including angiogenesis, DNA damage response, vesicle trafficking, extra-cellular matrix remodeling, protein homeostasis, and metabolic regulation. Integration of these transcript-level features with clinical variables improved risk stratification compared with clinical variables alone, indicating that isoform-level measurements capture multidimensional aspects of tumor aggressiveness that are not reflected by gene-level data alone. Although external validation in the CPTAC-ccRCC cohort was limited by isoform availability and therefore relied on differential isoform expression rather than transcript usage modeling, higher expression of *FGFR1:ENST00000326324* and *NUCB1:ENST00000405315* remained associated with worse survival. This finding supports the potential prognostic relevance of isoform-level features across independent cohorts. Notably, *NUCB1* is not prognostic at the gene level in ccRCC, further highlighting the added value of isoform resolution^64^.

Taken together, this study supports a model in which alternative splicing and transcript usage act as dynamic regulatory mechanisms in ccRCC, shaping tumor behavior by modulating coding capacity, pathway output, and cellular adaptation across key processes, including epithelial dedifferentiation, lipid metabolism, immune signaling, cytoskeletal organization, and angiogenesis. These findings should be interpreted in the context of limitations inherent to transcript-level analysis, including sensitivity to RNA quality and incomplete representation of splicing regulation by epigenetic and post-transcriptional mechanisms. In addition, constraints of the compositional DTU framework and convergence challenges in covariate-rich models limited full adjustment for potential confounders. Nonetheless, our findings overall support the biological relevance of transcript-level remodeling.

## Acknowledgments/Funding Information

This work was primarily supported by the Congressionally Directed Medical Research Programs, U.S. Department of Defense (CDMRP/DoD), grant W81XWH-20-1-0778, and by the National Institute of General Medical Sciences (NIGMS) under award P20GM104416. Additional support was provided by grants P30CA023108, P20GM130454 (NIGMS); R01CA275974, and R01CA253976 (National Institute of Health). Also, support from the Shared Resources at Dartmouth was received: Genomics and Molecular Biology — 10x Chromium — single cell & spatial genomics:S10OD030242,P30CA023108,P20GM130454,S10OD025235, RRID:SCR_021293

## Genarative

*AI (ChatGPT 5.5) and Grammarly services were used to assist with the preparation of this manuscript in the context of reviewing grammar and organizing the reading flow of the text*.

## Data Availability Statement

The data generated in this study are available upon request from the corresponding author.

## Code Availability Statement

R Script and Developed code for all analyses in this study are linked here: https://github.com/SalasLab/DTU-Analysis-in-ccRCC-Reveals-Isoform-Level-Remodeling

## Supplementary data

### Supplementary Tables

Table S1A: Increased-usage differentially used transcripts (DTUs) identified between ccRCC tumors and adjacent normal tissues. Table S1B: Decreased-usage differentially used transcripts (DTUs) identified between ccRCC tumors and adjacent normal tissues. Table S1C: Differentially used transcripts exhibiting discordant directionality relative to gene-level differential expression. Table S1D: Sensitivity analysis of increased-usage DTUs following exclusion of low-count samples. Table S1E: Sensitivity analysis of decreased-usage DTUs following exclusion of low-count samples.

Table S2A: HR and 95% CI for Cox-ph adjusted models for survival analysis between clusters and AUA Risk (Binary)

Table S3A: Increased-usage differentially used transcripts (DTUs) identified between ccRCC tumor clusters (C1 vs C2). Table S3B: Decreased-usage differentially used transcripts (DTUs) identified between ccRCC tumor clusters. Table S4. PFAM Domain annotations for the top 5 DTUs between tumor clusters.

### Supplementary Figures

Figure S1. Functional enrichment of genes with differential transcript usage in ccRCC. Figure S2. Representative transcript usage patterns and isoform switches in ccRCC tumors and adjacent normal kidney tissues. Figure S3: Figure S3. Differential expression and functional over-representation analysis of functional and non-functional coding transcripts in ccRCC Figure S4: Figure S4. Gene Ontology over-representation analysis of transcript classes differentially expressed between tumor cluster 2 and cluster 1. Figure S5: Figure S5. Performance of transcript-based prognostic models across resampling iterations.

## Abbreviations and Terminology

### Abbreviations

AF: alternative first exon
AUA: American Urological Association
A3: alternative 3′ splice site
A5: alternative 5′ splice site
BAM: binary alignment map
bp: base pairs
ccRCC: clear cell renal cell carcinoma
CDF: cumulative distribution function
CI: confidence interval
CPM: counts per million
CPTAC: Clinical Proteomic Tumor Analysis Consortium
DCC: Dartmouth Cancer Center
DEG: differentially expressed gene
DGE: differential gene expression
DNA: deoxyribonucleic acid
DRIMSeq: Dirichlet-multinomial framework for differential transcript usage analysis
DTU: differential transcript usage
EMT: epithelial-to-mesenchymal transition
ENmix: ENmix methylation preprocessing pipeline
ERV: endogenous retrovirus
ES: exon skipping
FDR: false discovery rate
FGFR1: fibroblast growth factor receptor 1
FPKM: fragments per kilobase per million mapped reads
GO: Gene Ontology
GSEA: gene set enrichment analysis
HiTIMED: hierarchical tumor immune microenvironment epigenetic deconvolution
HiTAIC: Hierarchical Tumor Artificial Intelligence Classifier
HIF: hypoxia-inducible factor
HR: hazard ratio
IRB: institutional review board
LASSO: least absolute shrinkage and selection operator
lncRNA: long non-coding RNA
logFC: log2 fold change
mRNA: messenger RNA
MX: mutually exclusive exons
NMD: nonsense-mediated decay
NMF: non-negative matrix factorization
OS: overall survival
PCA: principal component analysis
PCR: polymerase chain reaction
PD-1: programmed cell death protein 1
PI3K: phosphoinositide 3-kinase
QC: quality control
RCC: renal cell carcinoma
RI: retained intron
RIN: RNA integrity number
RNA: ribonucleic acid
RNA-seq: RNA sequencing
SE: skipped exon
STAR: Spliced Transcripts Alignment to a Reference
TCGA: The Cancer Genome Atlas
TEC: to be experimentally confirmed
TGF-β: transforming growth factor beta
TME: tumor microenvironment
TNM: tumor-node-metastasis
TPM: transcripts per million
VEGF: vascular endothelial growth factor
VHL: von Hippel–Lindau

## Authors contributions

- Conceptualization: CFN, LAS
- Data curation: CFN, YL, MZ
- Formal Analysis: CFN, YL,
- Funding acquisition: LAS, BCC
- Investigation: CFN
- Methodology: CFN, EP, FK, TJP, LPLAS
- Project administration: CFN, LAS
- Resources: CFN, LAS, FK, BR, JRP
- Software: CFN
- Supervision: LAS, BCC, PAP, LS
- Validation: CFN, EM
- Visualization: CFN, MZ
- Writing – original draft: CFN
- Writing – review & editing: CFN, JB, LAS, BCC, JDS, JRP, MSS

