## Supplementary Figures for "Differential Transcript Usage Reveals Isoform-Level Remodeling of Tumor Biology in Clear Cell Renal Cell Carcinoma"

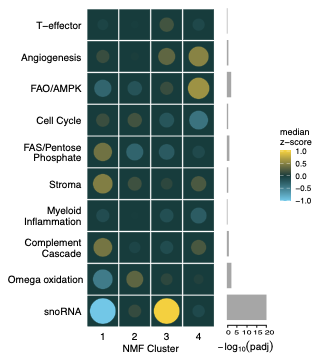

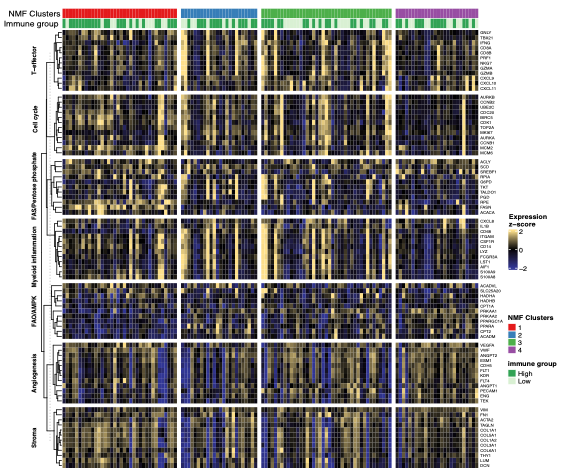

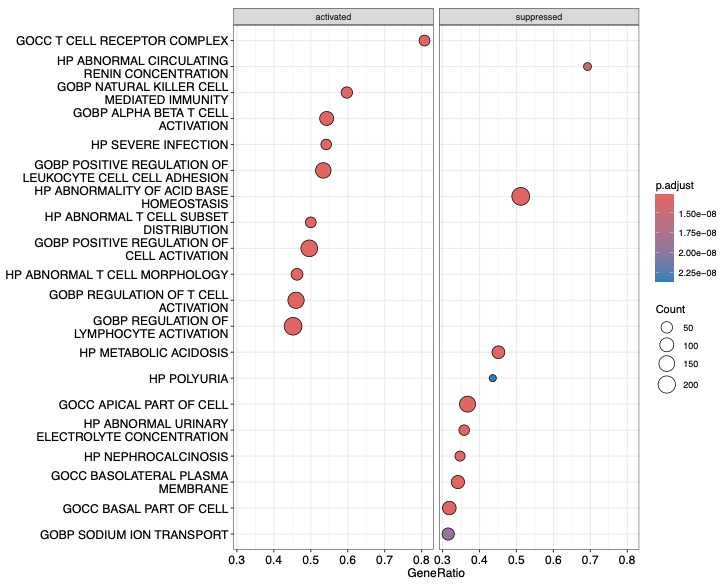

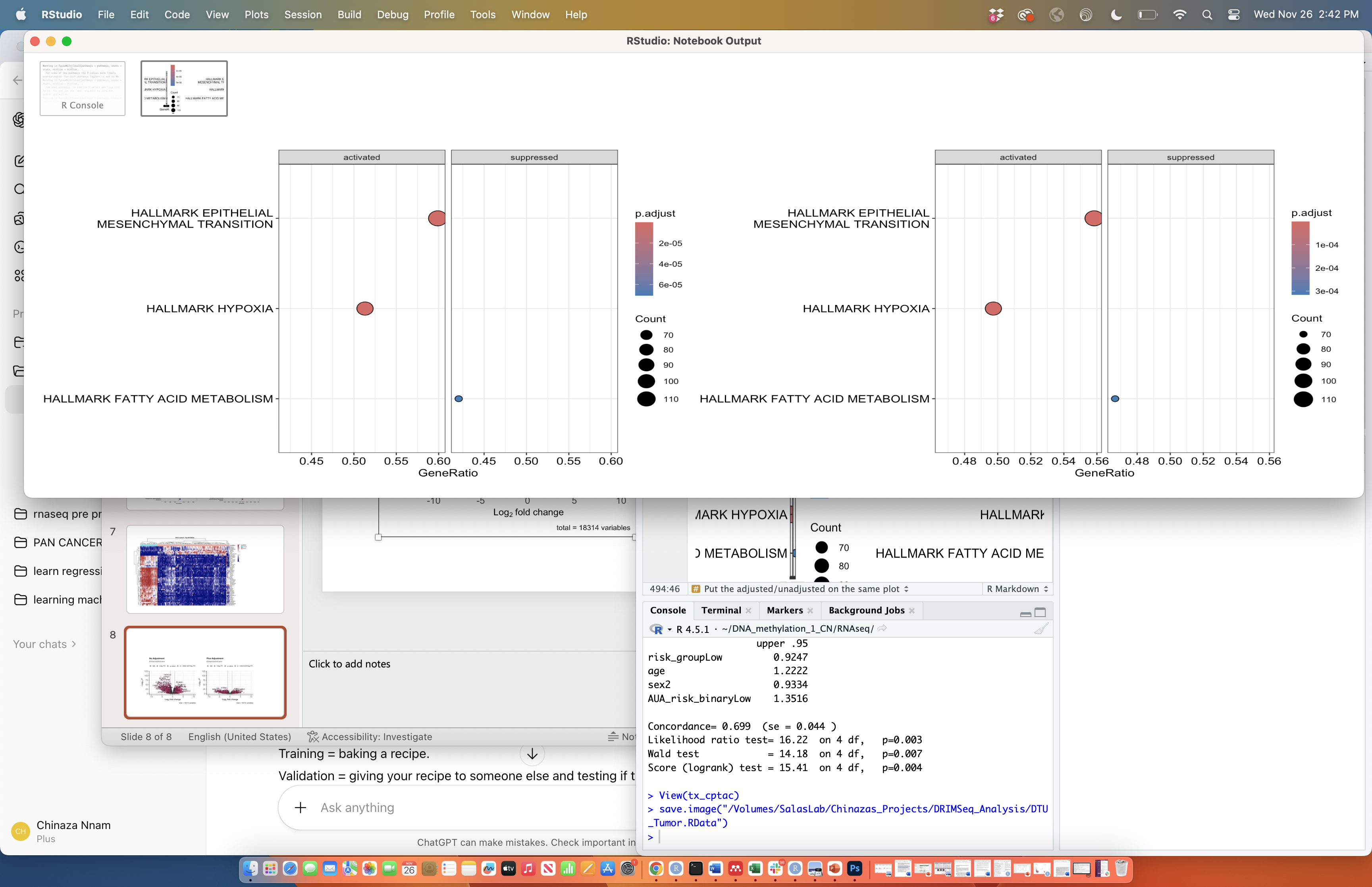

**D**

**C**

**B**

**A**

DEGs

gDTUs

987

1007

500

488

138

44

Total common =44+138= 182

**E**

**Figure S1. Functional enrichment of genes with differential transcript usage in ccRCC.
(A) Enrichment of selected MSigDB Hallmark terms among genes with differential transcript usage, showing activation of epithelial mesenchymal transition and hypoxia-related programs and suppression of fatty acid metabolism. (B) Enrichment of selected Hallmark C5 Category terms, showing activation of immune-related pathways and suppression of renal epithelial and solute transport associated processes. In both panels, dot size indicates gene count and color indicates adjusted *P* value. (C-D) Non-negative matrix factorization (NMF)-based clustering of DCC ccRCC tumors using IMmotion-derived molecular programs. Consensus clustering metrics, including the cophenetic correlation coefficient, residual sum of squares, and cluster stability profiles, supported selection of k = 4 as the optimal number of clusters. Bubble plot displays median z-scored enrichment of molecular programs across the four NMF clusters. Dot size represents −log10(adjusted P value), and color intensity represents median z-score enrichment. Cluster 4 demonstrated prominent enrichment of angiogenesis and FAO/AMPK-related metabolic programs. (E) Overlap between DEGs and gDTUs in ccRCC tumors versus adjacent normal tissues. Red indicates genes/transcripts with increased expression or usage in tumors, while blue indicates decreased expression or usage. Shared regions represent genes identified by both DEG and DTU analyses.**

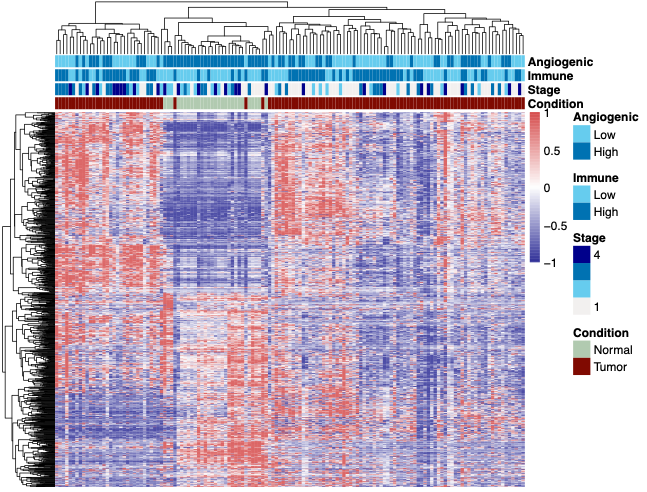

**A**

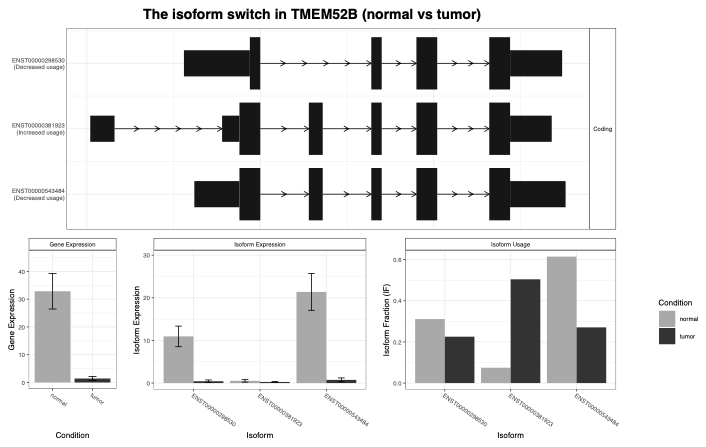

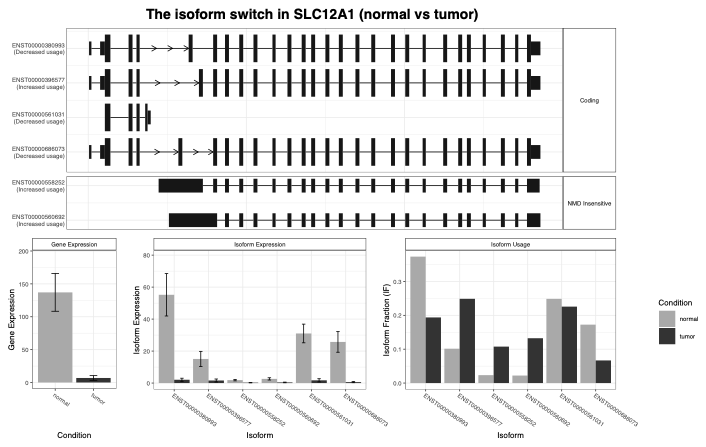

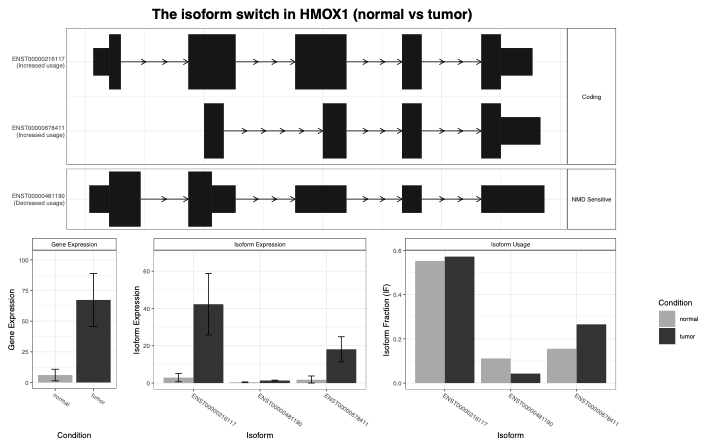

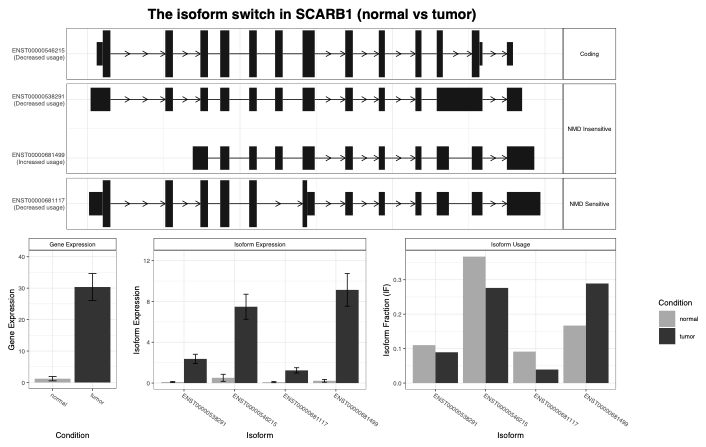

**Condition**

Normal

Tumor

**B**

**D**

**C**

**E**

**Figure S2. Representative transcript usage patterns and isoform switches in ccRCC tumors and adjacent normal kidney tissues.
(A) Heatmap of significant differentially used transcripts(n=1170) across normal kidney and ccRCC tumor samples, annotated by condition, stage, immune score, and angiogenic score.
(B-E) Isoform switch plots for HMOX1 and SCARB1, the top two transcripts with increased usage in tumors, and for SLC12A1 and TMEM52B, representative transcripts with decreased usage in tumors. Transcript structures are shown above bar plots of gene expression, isoform expression, and isoform usage, with gray indicating normal and black indicating tumor.**

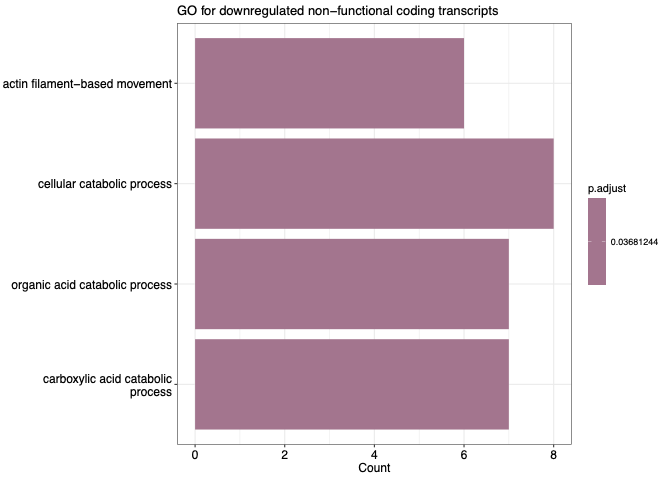

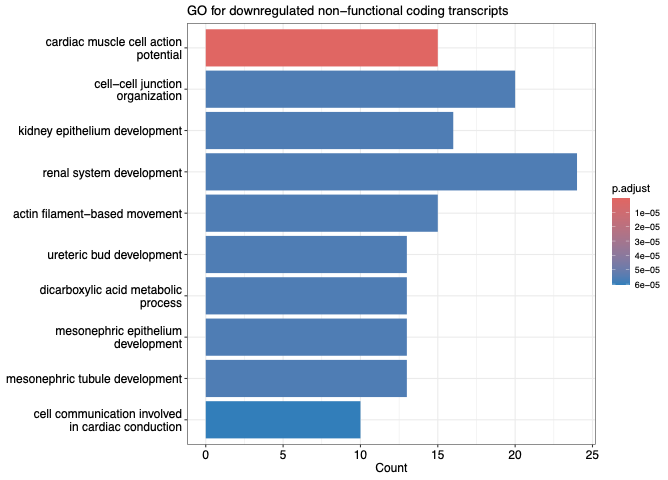

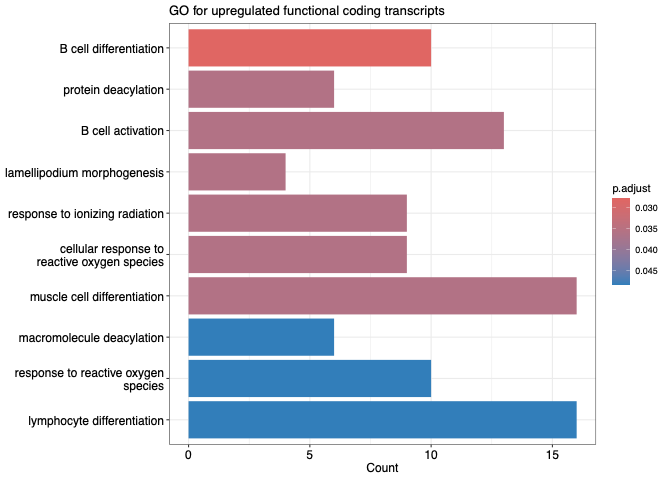

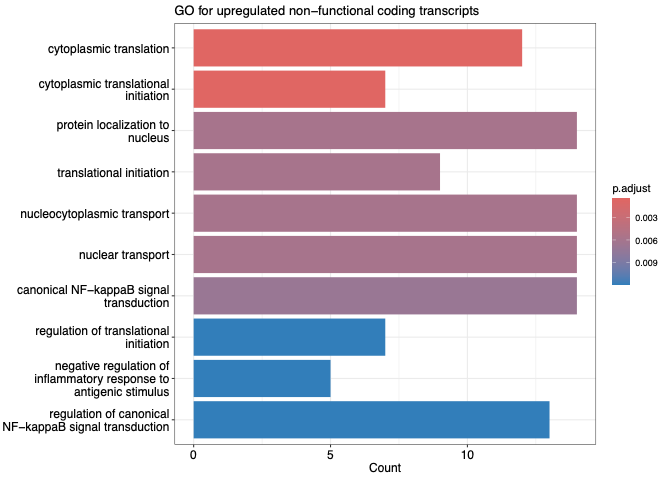

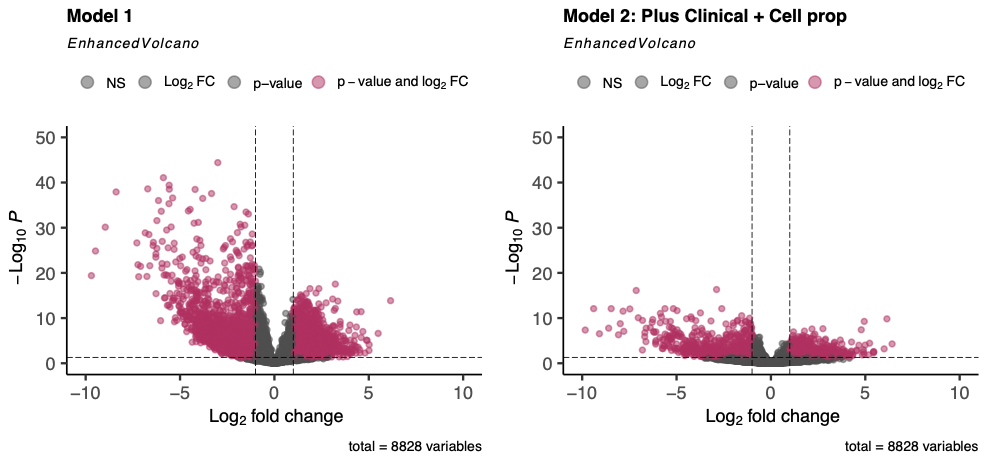

Baseline Model: +age, sex

Cell Type Adjusted Model: +age, sex, immune, angiogenic cell proportions

**A**

**E**

**D**

**C**

**B**

**Figure S3. DTU between tumor and adjacent normal samples and over-representation analysis of functional and non-functional coding transcripts in ccRCC.
(A) Enhanced volcano plots showing DTU from the baseline model and the model adjusted for clinical covariates and cell proportion.
(B-E) Gene Ontology over-representation analyses of upregulated and downregulated functional coding and non-functional coding transcripts. Bar length indicates transcript count, and bar color indicates adjusted *P* value.**

**Figure S4. Gene Ontology over-representation analysis of transcript classes differentially expressed between tumor cluster 2 and cluster 1.
(A,B) Enriched terms among upregulated and downregulated functional coding transcripts in cluster 2 relative to cluster 1.
(C,D) Enriched terms among upregulated and downregulated non-functional coding transcripts in cluster 2 relative to cluster 1. Bar length indicates transcript count, and bar color indicates adjusted *P* value.**

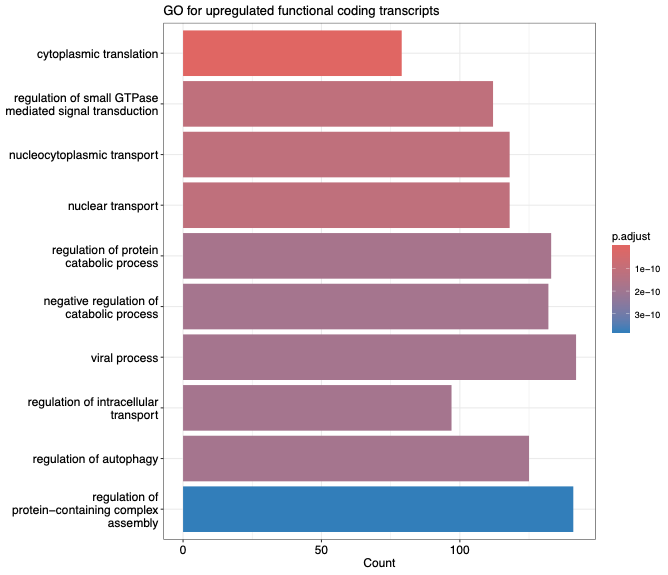

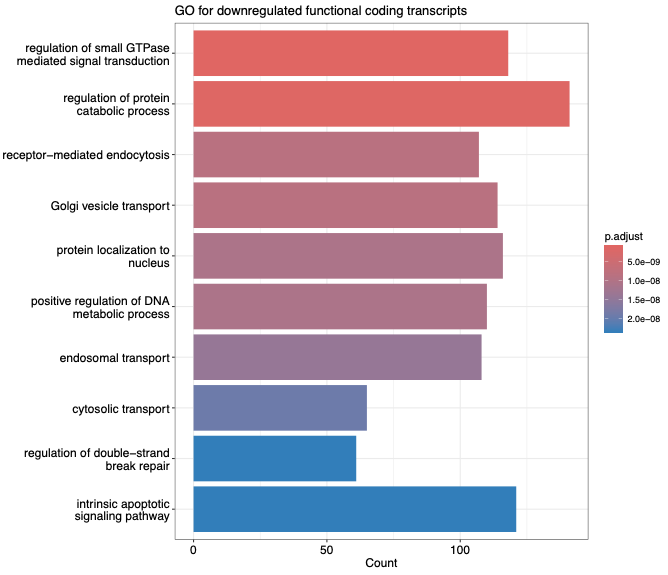

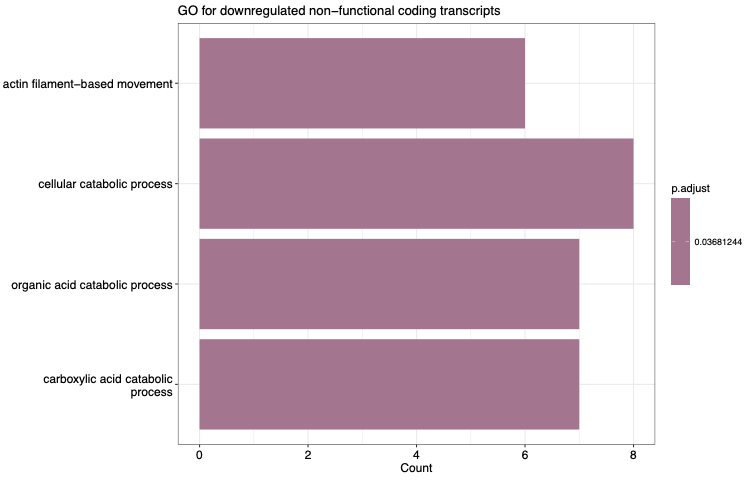

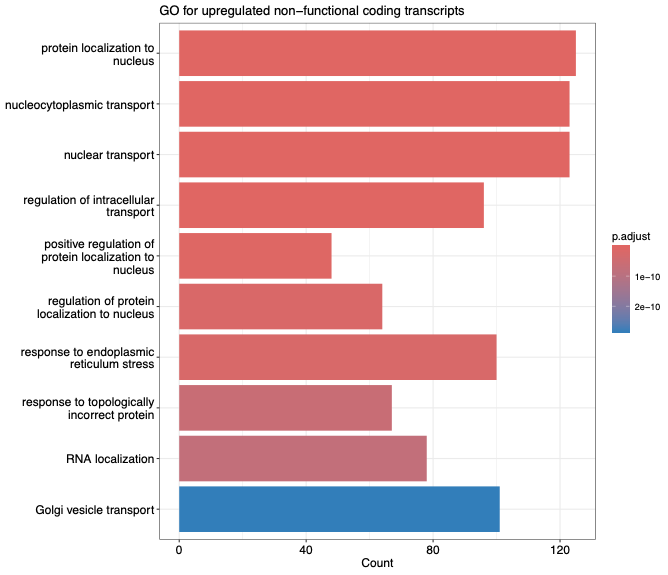

**A**

**B**

**D**

**C**

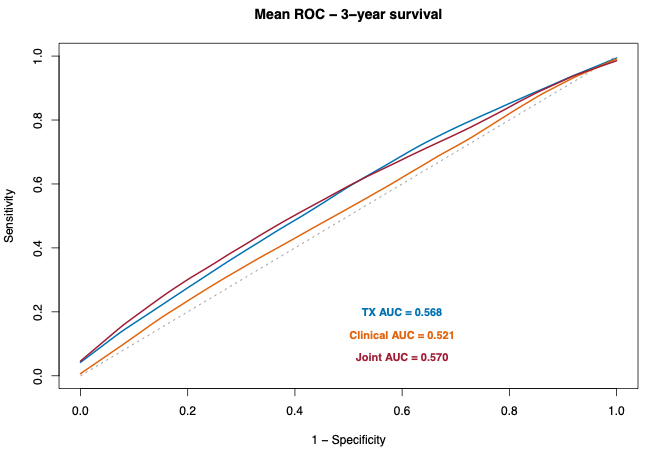

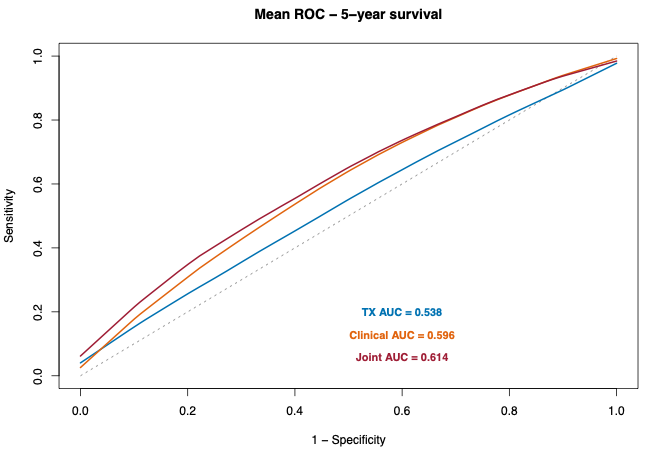

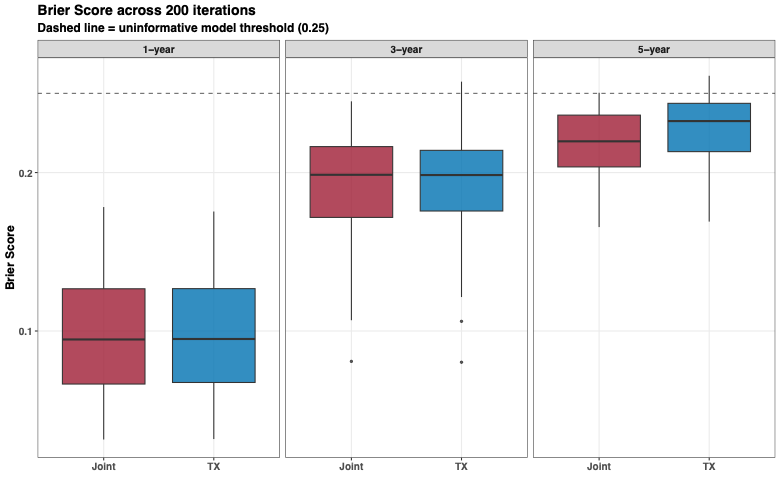

**A**

**B**

**C**

**Figure S5. Performance of transcript-based prognostic models across resampling iterations.
(A–B) Mean receiver operating characteristic (ROC) curves for 3-year (A) and 5-year (B) overall survival, averaged across 200 bootstrap iterations. Curves are shown for transcript-only (TX), clinical-only, and joint (combined transcript and clinical) models, with corresponding mean AUC values indicated.
(C) Brier scores across 200 bootstrap iterations for 1-year, 3-year, and 5-year survival predictions, comparing transcript-only (TX) and joint models. Lower Brier scores indicate improved predictive accuracy. The dashed line represents the uniform model threshold (0.25).**
